# Communication through autoattractants can enhance and limit collective migration of immune cells

**DOI:** 10.64898/2026.04.07.716888

**Authors:** David M. Versluis, Robert H. Insall

## Abstract

Many eukaryotic cells produce attractant molecules to which they themselves are also attracted. For example, neutrophils produce leukotriene B4 while swarming. These autoattractants create a secondary signalling layer that coordinates collective cell behaviour. The principles by which autoattractants shape migration, however, remain poorly understood.

Here we use a hybrid agent-based computational model to dissect the effects of autoattractants on the collective chemotaxis of immune cells. We find that autoattractant signals strongly enhance cells’ responses to primary attractant, but only if the autoattractant is sufficiently short-lived: at longer lifetimes, accumulation degrades the directional information cells can extract. The overall effect of autoattractants therefore depends crucially on the balance between production and efficient removal — whether through cellular uptake, enzymatic breakdown, or inherent chemical instability. Optimal lifetimes exist, determined by cell speed and attractant diffusion, yet remarkably independent of cell density and primary attractant concentration. Autoattractants whose removal is governed by inherent instability rather than direct cellular breakdown coordinate migration less efficiently, but work more robustly across different environments. In the absence of cell-mediated breakdown, the model further reveals a characteristic optimal cell–cell distance: too little communication leaves cells uncoordinated, while excessive signalling drives them into slow-moving aggregates. Strikingly, the conditions that produce the most efficient chemotaxis lie close to those that trigger aggregation, suggesting that many autoattractant systems operate near a critical boundary that can be tuned by evolution either to disperse cells or to bring them together.

**Author Summary:** Immune cells must travel through the body to reach sites of injury, infection, and developing tissue. They are guided by chemical signals. Many immune cells, such as neutrophils gathering at wounds and microglia in the brain, also release their own signals that attract others of the same kind. We call these ’autoattractants’. In effect, the cells tell each other where to go. This communication helps coordinate group movement, but little is known about how it works.

We built a computer simulation of migrating immune cells to explore how autoattractants shape their collective behaviour. We find that autoattractants must be short-lived to be useful: when autoattractant accumulates, it obscures any sense of direction. Signals that cells actively break down, or that decay on their own, carry clear directional information. With signals that decay on their own, a delicate balance emerges: just enough cooperation leads to the most efficient coordinated migration, while a little too much causes cells to clump together instead of moving forward. Our work suggests that many more cell types may use this communication than we know, and that the same system can be tuned by evolution to spread cells out or to gather them together.

## 1 Introduction

Many eukaryotic cells produce attractant molecules that they themselves are also attracted to. We refer to these molecules as “autoattractants”. There are many examples of this behaviour, in particular in mammalian immune cells. For example, neutrophils that sense the attractant peptide fMLP produce leukotriene B_4_ (LTB_4_), which functions as a signal-relay molecule during chemotaxis, amplifying a recruitment of additional neutrophils to sites of inflammation often characterised as ’swarming’ [1, 24, 30]. LTB4 is also quickly bound and sequestered from the medium by neutrophils [7]. In response to LPS neutrophils also produce IL-8, which functions as a chemoattractant for neutrophils [3, 4]. Autoattraction can also exist without swarming, however. For example, microglia are attracted to ATP, and also release it by exocytosis in response to LPS [10, 17], but they do not move in swarms as neutrophils do. This may be because ATP is degraded both near microglia and in the broader extracellular space [31, 8], which would limit swarm formation. Autoattractants can also be produced in response to attractant chemokines, such as in eosinophils, which produce the short-lived autoattractant prostaglandin D2 in response to the activation of its chemokine receptor CCR3 [16, 2, 22, 28, 14].

Despite the prevalence of autoattractant signalling across these systems, the mechanics of transmitting information through autoattractants remain understudied. In particular, we focus on a specific class of interactions: the exchange of signals among identical cells during chemotaxis along self-generated gradients. Self-generated gradients are widespread among many cell types and attractants, and often play a large role in cell migrations [33, 32, 18]. We focus on self-generated gradients specifically because they are inherently temporary and dynamic. If a stable, externally imposed gradient were already present and sufficient for directed migration, the additional information carried by autoattractant signalling would be less useful.

As self-generated gradients are inherently temporary, they are hard to study *in vivo* and *in vitro*. We used computational modelling to create an overview of how cells, with a particular focus on immune cells, may use autoattractant signals to enhance collective chemotaxis in self-generated gradients. We based our modelling on earlier successful work using agent-based modelling of this system [32, 11], and we have previously used this technique to create predictions for migrating macrophages [20]. Crucially, we have expanded the conceptual framework to include autoattractants as separate species that can be produced, broken down, or decay in various ways. Here, we used modelling to explore: (i) how autoattractants can enhance the ability of cells to gather information during chemotaxis to weak or noisy signals; (ii) the trade-offs of various strategies for creating useful autoattractant signals; (iii) the interaction between cell speed and effective communication; and (iv) the conditions under which autoattractant signalling leads to aggregation rather than enhanced migration.

## 2 Results

### 2.1 Approach

We used a hybrid agent-based modelling approach in which cells are represented as individual off-lattice agents operating on a regular lattice of attractant concentrations. The model simulates a chemotaxis chamber: cells start at one end of a rectangular domain filled with an attractant that is initially present, which we call the primary attractant. The model is then run in discrete timesteps to generate predictions for cell behaviour and attractant concentrations across the environment over time. This approach can represent many different types of migrating immune cells and situations. We will describe the working of the model here briefly, and refer to the methods for a full description.

Each cell is approximated by a circle (Fig. 1A), representing the area over which a cell can sense, add, and remove attractants within each simulation timestep (6 s). Each timestep, the cell uses saturable receptors to sense and break down primary attractant in the area it covers. It then moves in the direction of the gradient it senses (Fig. 1B). Random noise is added to cell perception to prevent excessively shallow gradients from being followed, and the attractant diffuses between timesteps so that smooth gradients can be formed. The combination of attractant removal and movement towards attractant leads to a self-generated gradient and limited migration away from high densities of cells (Fig. 1C). We have previously shown that this approach leads to a good reproduction of migration to self-generated gradients [11, 33].

**Figure 1:**
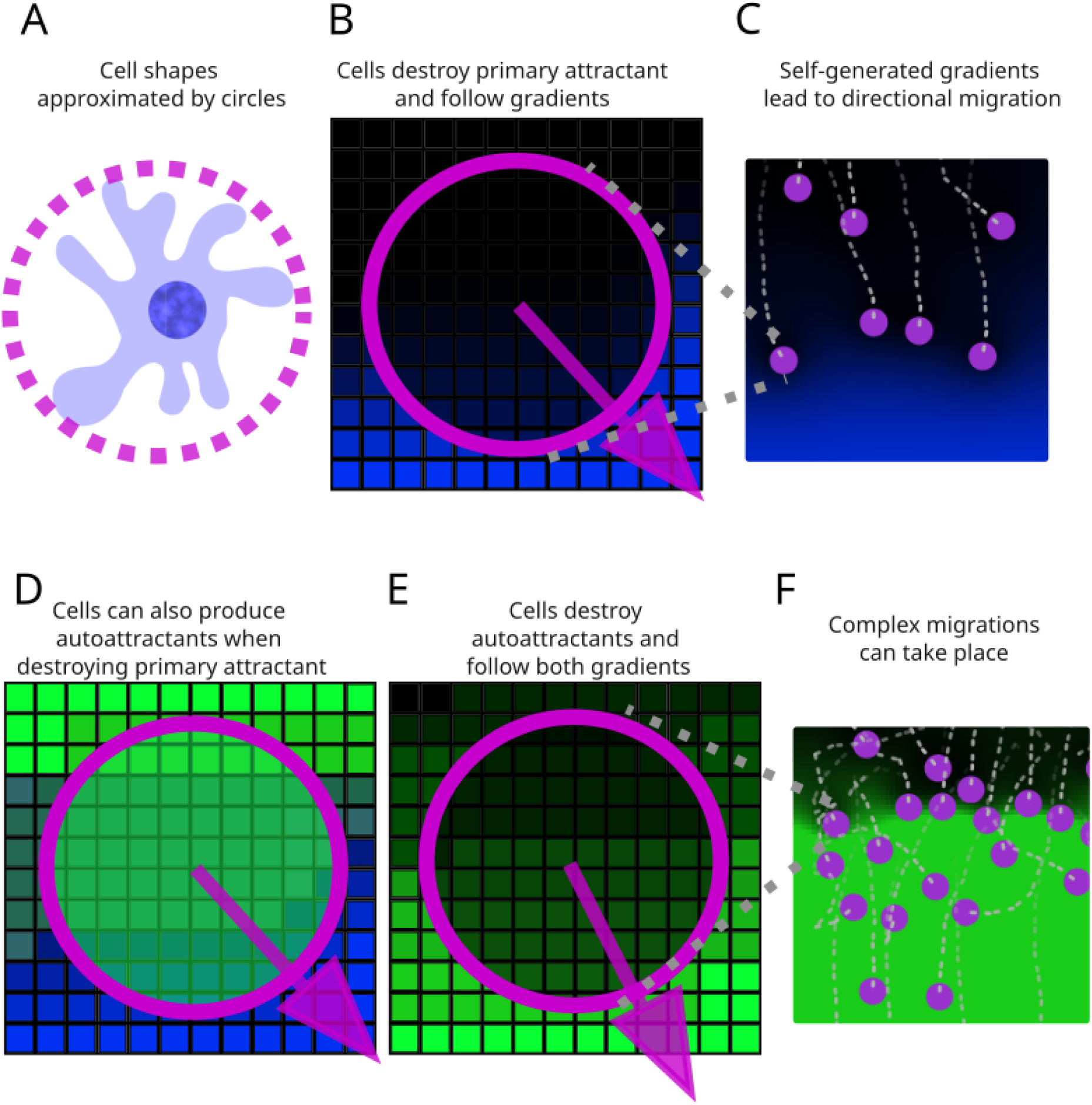
Schematic model setup. **A.** The area within which a cell can sense attractants is approximated by a circle. **B.** Schematic of a single modelled cell breaking down primary attractant while also following its gradient. **C.** Visualisation of cells chemotaxing to a self-generated gradient of primary attractant in a sample simulation. **D.** Schematic of cell producing autoattractant while creating and following a gradient mostly composed of primary attractant. **E.** Schematic of a cell breaking down autoattractant and following a gradient mostly composed of autoattractant. **F.** Visualisation of cells chemotaxing to a self-generated gradient of autoattractant in a sample simulation.

Cells also produce autoattractant, which is modelled similarly to the primary attractant, except that it is not initially present but instead produced by the cells in proportion to the amount of primary attractant they break down (Fig. 1D). For example, if a cell breaks down 1 unit of attractant in a timestep and the ratio is set to 10, the cell will produce 10 units of autoattractant. The ratio of production is determined by the parameter settings of specific simulations. Cells also sense and break down autoattractant using receptors separate from those used for primary attractant (Fig. 1E). Each cell uses information from both kinds of receptor to arrive at a single target direction, towards which it will move with a fixed speed. Though speed per model timestep is fixed, cells will vary in how consistently they head towards the migration goal, so the speed measured over timescales longer than one timestep (6 seconds) will also vary. Taken together, this leads to complex migrations, where cells can indirectly follow the gradient of primary attractant through following the autoattractant that was produced based on it (Fig. 1F). We used several sets of parameters for the model, based as far as possible on plausible values for immune cells (Table 1 and 2).

**Table 1:**
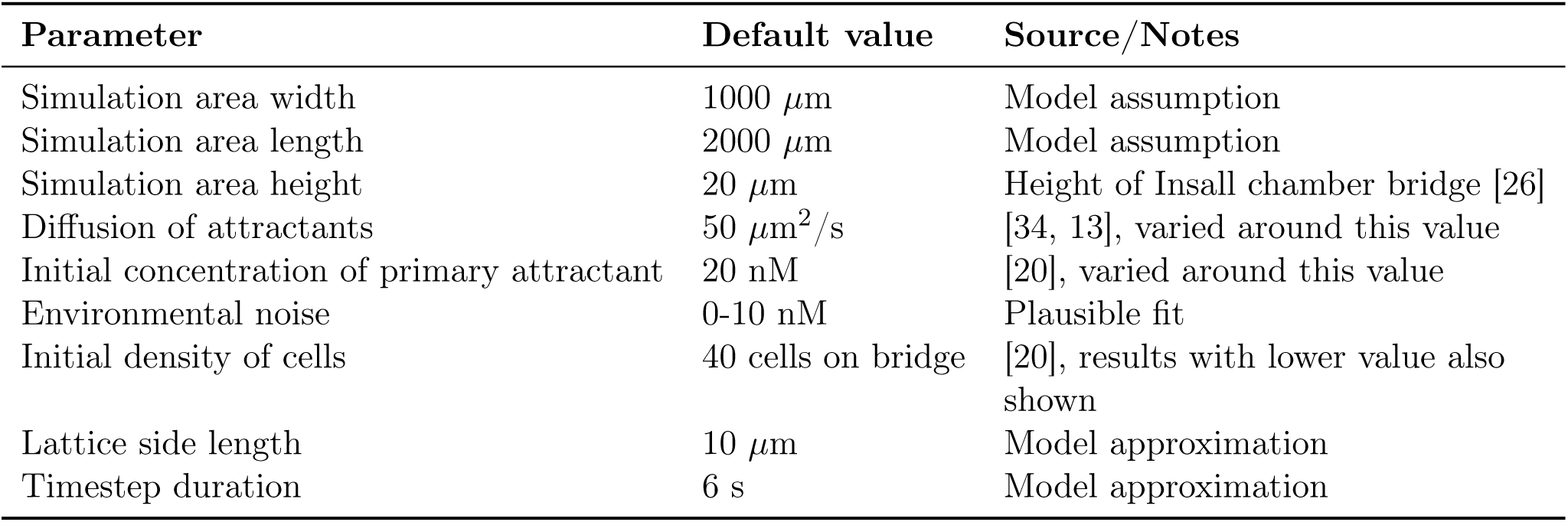
Environmental and simulation parameters.

### 2.2 Autoattractants can improve migration

We first initialised the model with a parameter set based on fast moving immune cells (Table 1 and 2), such as neutrophils. We initialise all cells on one side of the rectangular area, and consider a cell to have migrated successfully if it is in the bottom quarter of the area by the end of the simulation. As a control, we first initialise the model with no autoattractant production. The primary attractant on its own can only recruit a limited number of cells (Fig. 2A, Video S1). When we add autoattractant production by cells we find that more cells migrate successfully (Fig. 2B, Video S1) and a clear gradient of autoattractant is formed (Fig. 2B, right panel).

**Figure 2:**
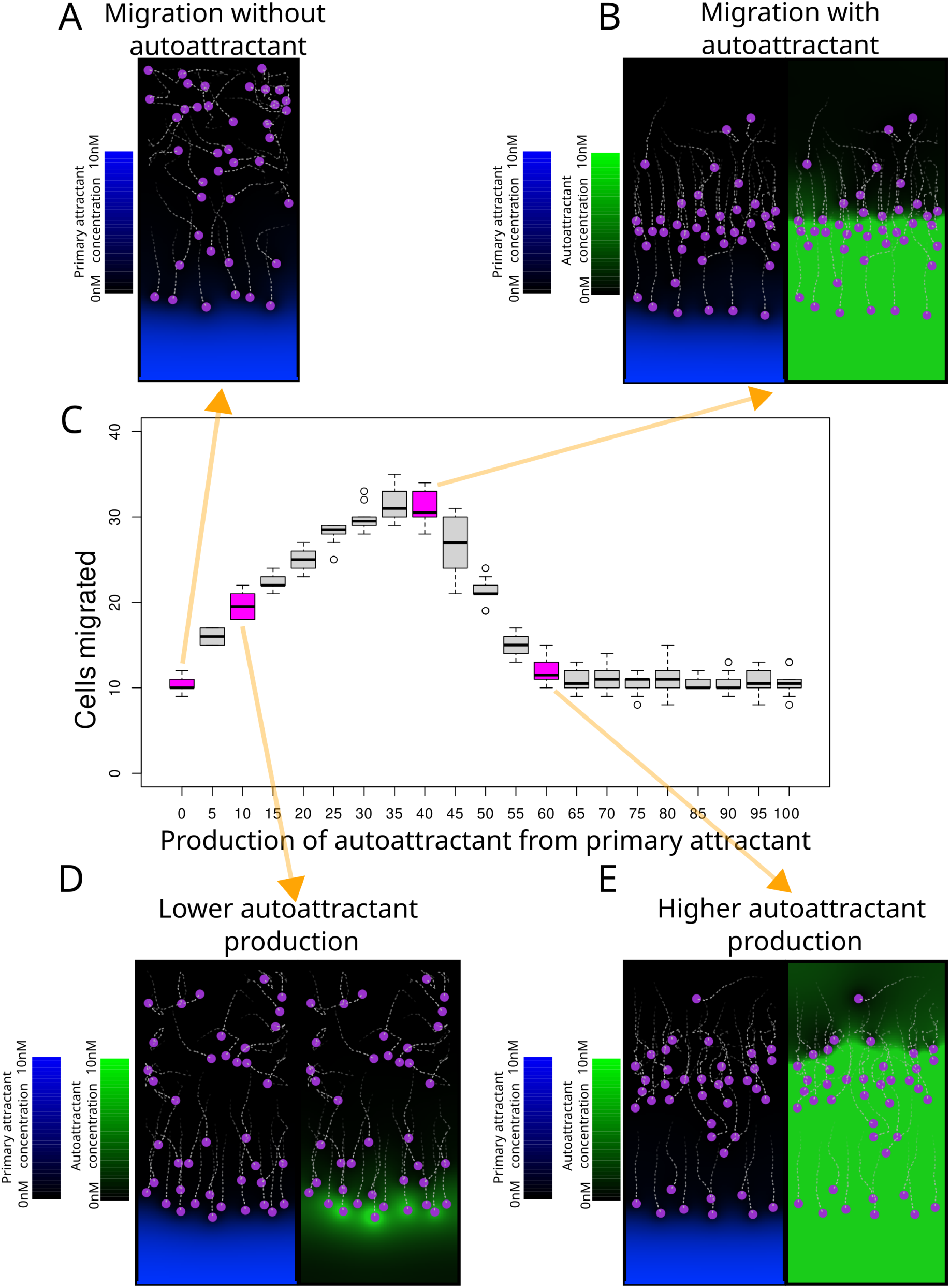
Autoattractant leads to more migration at correct production ratios. **A.** Cell locations and primary attractant distribution during a representative model simulation with only a primary attractant. **B.** Cell locations and primary attractant (left) and autoattractant (right) concentration during a model simulation with an added autoattractant. **C.** Cell migration over a wide range of autoattractant production ratios. *n* = 10 per condition. **D,E.** Visualisations as in A of the effects of lower and higher production, which both lead to less migration compared to B, as either a very small autoattractant gradient forms (D), or it forms very far away (E).

The number of cells that migrate depends on the amount of autoattractant produced (Fig. 2C), but with a clear optimal level above which migration returns to a baseline level similar to that without autoattractant production. Interestingly, no amount of autoattractant causes less migration than that of the baseline without autoattractant. Smaller amounts than the optimum lead to a smaller migration (Fig. 2D, Video S1), as the wave of autoattractant formed by the frontrunners is broken down before it reaches all cells. Larger amounts also lead to less migration, as the part of the autoattractant gradient that can be sensed is further away from the first wave (Fig. 2E, Video S1). The front wave still functions normally: as the concentration is very high, the autoattractant receptors are saturated in all directions.

### 2.3 Optimal autoattractant production and breakdown depend on environment, but optimal lifetimes are similar

To be effective, autoattractant must reach other cells that do not sense the primary attractant gradient, and do so in the right concentration. We have shown that there is a range of different autoattractant production rates that function, but we also expect the rate of breakdown to influence the effectiveness. By altering the maximum rate of autoattractant breakdown (Eq. 10) we can effectively vary the dynamics. We systematically analyse pairs of production and breakdown values across a wide range (Fig. 3A). This shows that there is a linear relationship between optimal production and optimal breakdown, leading to an optimal balance in parameter space (schematically shown in Fig. 3B). Where there is relatively too much production, autoattractant accumulates as in Fig. 2E. Where there is relatively too much breakdown there is insufficient autoattractant to form a large wave, as in Fig. 2D.

**Figure 3:**
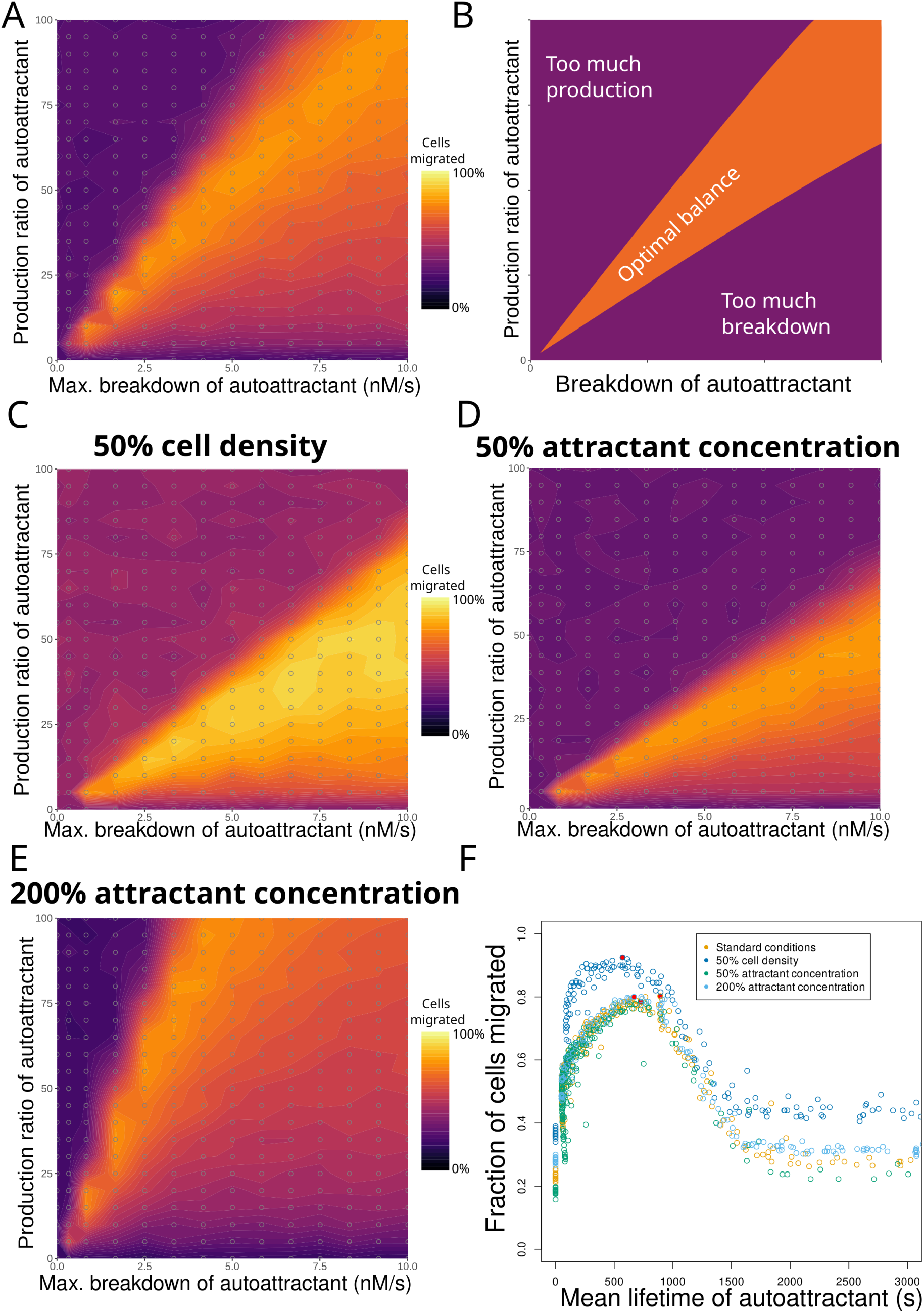
Optimal production and breakdown values vary, but optimal autoattractant lifetimes are similar. **A.** The effect of autoattractant production (as a ratio of breakdown of primary attractant) and autoattractant breakdown on migration. *n* = 10 per point. **B.** Schematic of the relation from A, highlighting that an optimal balance exists between the production and the breakdown of autoattractant. **C–E.** The combined effect of production and breakdown of autoattractant for different cell densities and attractant concentrations. *n* = 10 per point. **F.** Overview of the relation between migration and the mean lifetime of autoattractant for each parameter combination in A,C,D&E. Capped at a mean lifetime of 3000 s for readability. *n* = 10 per point. Filled circles represent the best migration for that condition.

We next investigated how robust this migration is by looking into how different initial conditions may influence this relationship. In an *in vivo* context cells cannot expect to always encounter the same concentration of attractant, or to be at a high density optimal for collective migration. We test a lower cell density, a higher initial concentration of primary attractant, and a lower initial concentration of primary attractant (Fig. 3C–E). These show different optimal production and breakdown balances. This means that the same cells may chemotax much more or less depending on these environmental factors. However, we can show that the dynamics at play are still the same by calculating the average time that an autoattractant molecule can be expected to exist, given how much autoattractant is produced and how much is broken down. We call this the lifetime of the autoattractant. In Fig. 3F we examine the relationship between this lifetime and migration. This shows that all simulations in which cells migrate efficiently are within the same range of autoattractant lifetimes, between roughly 250 and 1000 seconds, despite differences in the environmental factors.

### 2.4 Decaying autoattractants allow for migration less dependent on environment

We showed how similar autoattractant lifetimes are optimal across environments, despite requiring very different amounts of autoattractant production and breakdown. This suggests that an autoattractant with a fixed lifetime could cause a very consistent migration in different environments. This could occur if the autoattractant decays at a consistent rate, due to enzymes being present across the tissue or due to the autoattractant oxidising rapidly on its own. Examples are ATP [8] and prostaglandin D2 [22, 28, 14]. We implement this as an autoattractant that is continuously broken down over time everywhere, instead of only near cells, so that its decay depends only on time. Effectively, the autoattractant always has a fixed lifetime. Similar to the broken down autoattractant, this decaying autoattractant leads to much more effective migration than without an autoattractant (Fig. 4A, Video S2). As before, we systematically examine the relation between production and decay in this system, and a very different relationship emerges: above a certain production, the same decay is optimal (Fig. 4B, Video S2). There is now accumulation of autoattractant when decay is slow, leading to accumulation of autoattractant without a discernible gradient. An autoattractant that decays very quickly similarly does not lead to a discernible gradient, leading to poor performance at either extreme. The optimal values remain similar when we vary cell density and attractant concentration (Fig. 4C–E). We can compare how well the outcomes of different parameters correlate with the baseline by examining the relation between the number of cells migrated for the same combination of autoattractant production and breakdown or decay. This shows that for cells with a broken down autoattractant the same parameters cause very different levels of migration depending on the initial attractant concentration and cell density (Fig. 4F), but with a decaying autoattractant the same parameter values produce migrations that are well-correlated between conditions (Fig. 4G).

**Figure 4:**
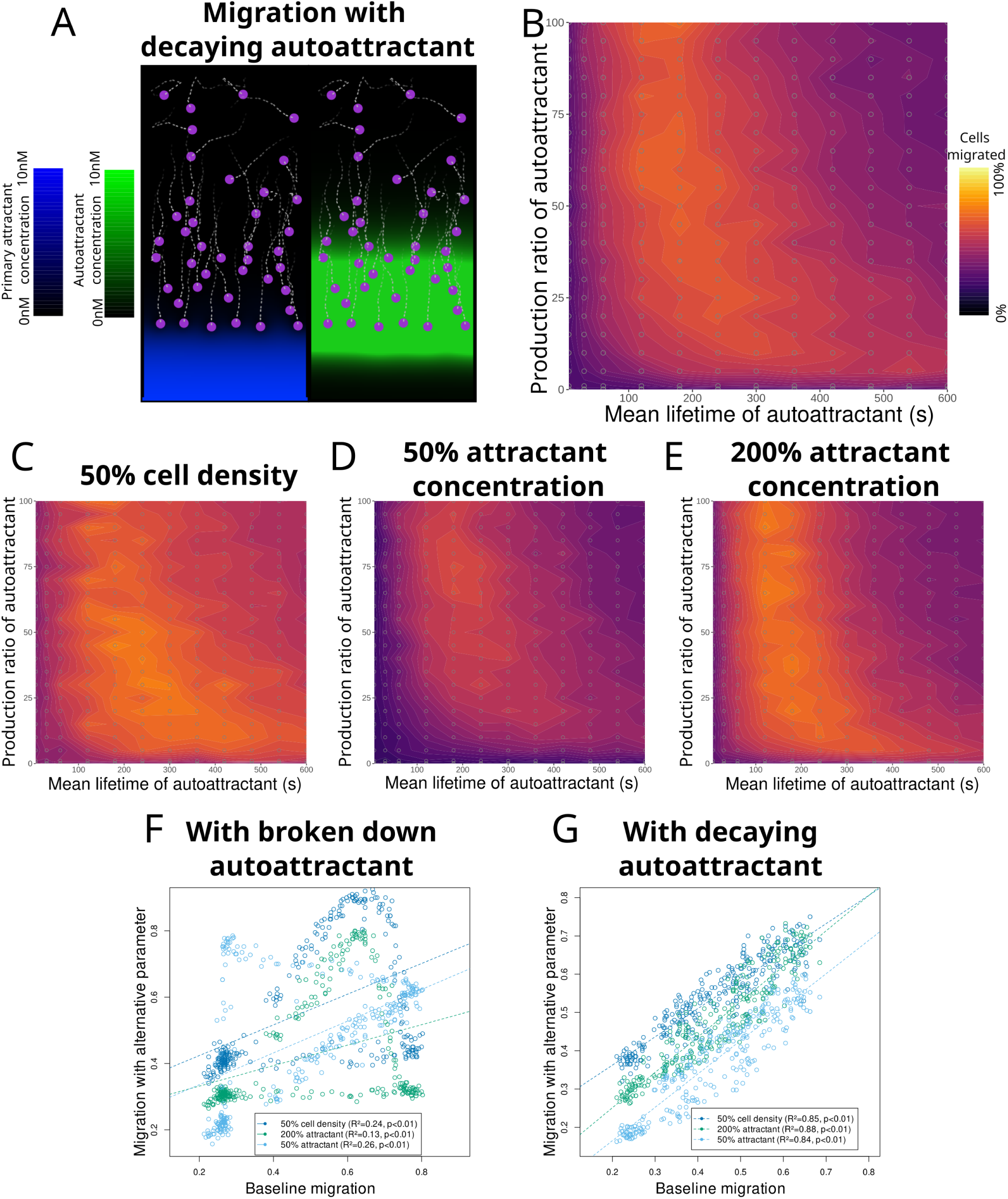
Decaying autoattractants lead to migration that is more stable across environments. **A.** Cell locations and primary attractant (left) and autoattractant (right) concentration in a sample model simulation with an autoattractant that decays constantly instead of being broken down by cells. **B.** The relationship between autoattractant decay and production and migration outcomes. *n* = 10 per point. **C–E.** As in B, but for low cell density, low initial primary attractant concentration, and high initial primary attractant concentration, respectively. **F.** The relationship between migration efficiency with the baseline conditions of Fig. 3A (on the X axis) compared to those of Fig. 3C, D, and E (on the Y axis), each dot represents one combination of autoattractant production and autoattractant breakdown. **G.** The relationship between migration efficiency with the baseline conditions of F_1_ig_1_. 4 B compared to those of 4 C,D, and E. Each dot represents one combination of autoattractant production and autoattractant breakdown, as in the previous panel.

### 2.5 Slow-moving cells can migrate better or aggregate

We have previously examined only fast-moving migrating cells, with parameters based on measurements of neutrophils, but many immune cells, such as macrophages or microglia, move much more slowly. We now examined how a parameter set with cells that move 1/20th of the speed of the cells we studied previously (Table 2) affects the migration. We assumed these cells also break down the primary attractant slower, so that they still create discernible self-generated gradients, and we give them an appropriate amount of time to reach the other side of the environment.

**Table 2:** Cell-specific parameters.

| Parameter | Fast-moving cells<br>(e.g. neutrophils) | Slow-moving cells<br>(e.g. macrophages) | Source/Notes |
| --- | --- | --- | --- |
| Simulation duration | 1000 steps (100min) | 20000 steps (2000min) | Plausible fit |
| Movement speed | 20 $\mu\text{m}/\text{min}$ | 1 $\mu\text{m}/\text{min}$ | [15, 20] |
| Minimum time to reverse direction | 2.8 min | 2.8 min | [19] |
| Sensing distance from cell centre | 30 $\mu\text{m}$ | 30 $\mu\text{m}$ | Plausible fit |
| Receptor $K_d$ | 10 nM | 10 nM | [29] |
| $K_m$ of primary attractant breakdown | 10 nM | 10 nM | Matched with Kd |
| $V_{\text{max}}$ of primary attractant breakdown | 200 nM/min | 10 nM/min | Plausible fit |
| $K_m$ of autoattractant breakdown | 10 nM | 10 nM | Matched with Kd |
| $V_{\text{max}}$ of autoattractant breakdown | variable | variable | Varied throughout |
| Autoattractant production ratio $\alpha$ | variable | variable | Varied throughout |

We found that with a broken down autoattractant the general dynamics of the slow-moving cells were similar to those of the fast-moving cells, as autoattractants enhanced migration (Fig. 5A&B, Video S3), and there was a similar linear relationship with breakdown (Fig. 5C & Fig. S1) and an optimal lifetime that was roughly 20 times as long, in proportion to the change in cell speed, and which was again similar for all conditions, though with greater variability than before (Fig. 5D, Fig S1 A-C). Furthermore, we found that a decaying autoattractant again led to similar optimal lifetimes regardless of conditions (Fig. 5E & Fig. S1 D-F), but there was also an unusual new behaviour: with a large production of a rapidly decaying autoattractant, cells formed transient clumps (Fig. 5E, inset lower left, Video S4). These These clumps consist of cells whose steering is more biased towards neighbouring cells than up the primary attractant gradient. Together they moved slower than individual cells, leading to an inhibited migration that is worse than that without autoattractants, which never occurred with the fast-moving cells. Other conditions within this set showed varying degrees of efficiency and direct cell-cell contact (Fig. 5E, insets, Video S4).

**Figure 5:**
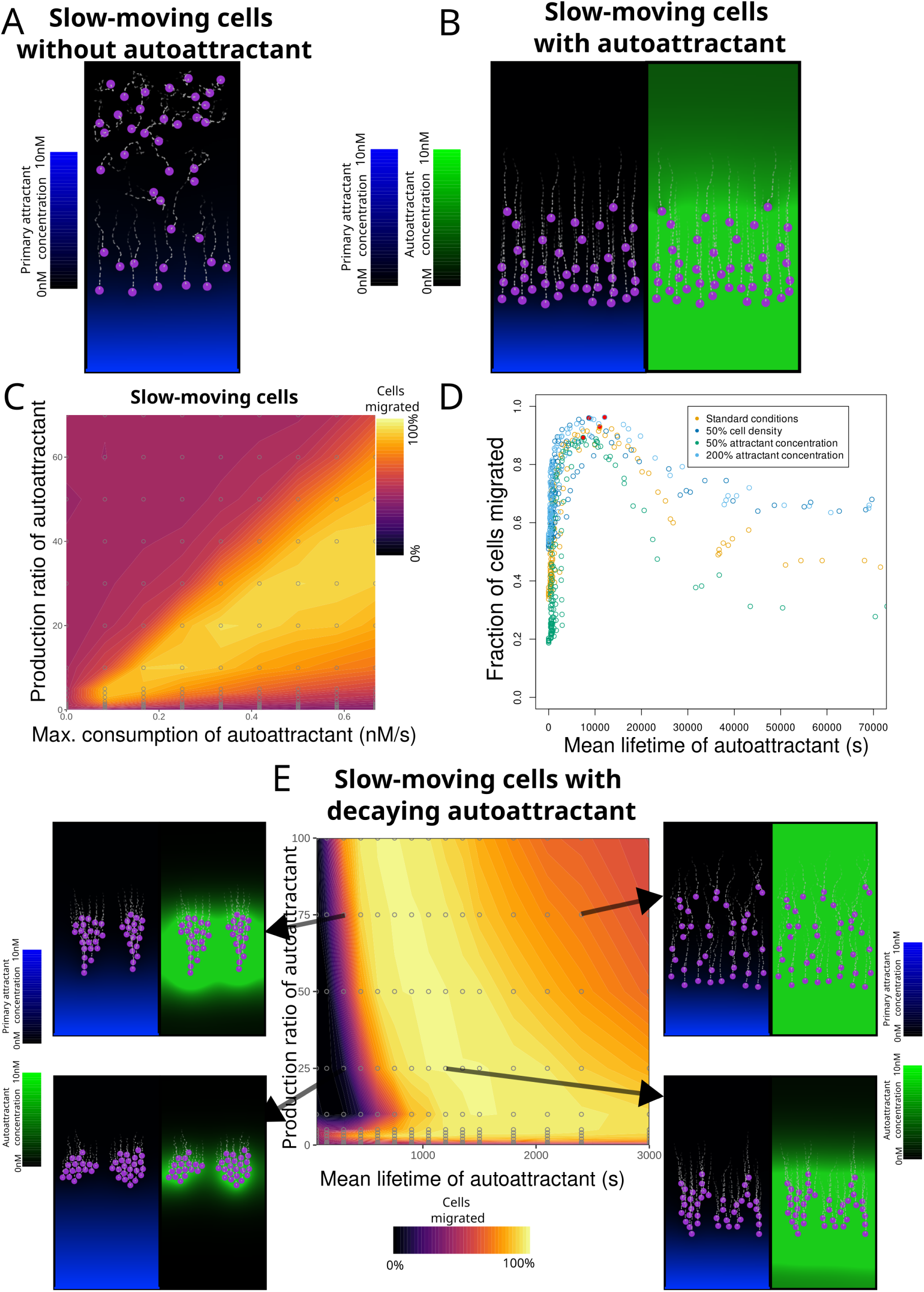
Slow-moving cells have longer optimal autoattractant lifetimes and can more easily aggregate during migration. **A.** Cell locations and primary attractant distribution in a representative model simulation with slower cells (Table 2) and only a primary attractant. **B.** As A, but with an added autoattractant that is broken down by the cells, displayed on the right. **C.** The relationship between autoattractant breakdown and production and migration outcomes. *n* = 10 per point. **D.** Overview of the relation between migration and the mean lifetime of autoattractant for the parameters in C, as well as with lower cell density and lower and higher initial primary attractant concentration. Capped at a mean lifetime of 70000 s for readability. *n* = 10 per point. Filled circles represent the best migration for that condition. **E.** The relationship between autoattractant decay and production and migration outcomes. *n* = 10 per point. Insets show examples of different behaviour across this spectrum.

### 2.6 Increased diffusion of attractant leads to shorter optimal lifetimes and increased clumping

We noted that the conditions where autoattractants inhibited effective migration were very close to those that promoted effective migration. To gain additional insights into the role of the attractant and autoattractant in this process we examined both slower and faster diffusion speeds for both attractants and autoattractants. Diffusion speeds of attractants vary across several orders of magnitude for different signalling molecules and conditions [34, 13], such as 6.2 *µ*m^2^/s for the chemokine CXCL13 [25] and up to 710 *µ*m^2^/s for ATP under ideal conditions [6]. We examined the effect of diffusion speeds 5x slower and 5x faster than our baseline (10 *µ*m^2^/s and 250 *µ*m^2^/s) with a broken down autoattractant for both fast (Fig. 6A & Fig. S2) and slow (Fig. 6B & Fig. S3) cells. We found that the optimal mean lifetime was shorter with faster autoattractant diffusion in both conditions, regardless of attractant diffusion speed. Slower autoattractant diffusion speeds consistently led to poorer migration with slow cells, but not fast cells. However, the shape of the relationship between production and breakdown was maintained throughout. We next examined how different diffusion speeds affect migration with decaying autoattractants. Fast-moving cells maintained similar dynamics regardless of diffusion speed, though migration was impaired in the more extreme cases (Fig. S4). For slow-moving cells, however, the effects depended on which substance diffused faster. Increasing attractant diffusion alone or decreasing autoattractant diffusion alone caused slow-moving clumps to form under many more parameter combinations (Fig. 6C, left middle and lower middle), as the short-range influence of the autoattractant increased relative to the attractant. Conversely, increasing autoattractant diffusion alone or decreasing attractant diffusion alone reduced the number of parameter combinations that led to slow-moving clumps (Fig. 6C, right middle, upper middle). When diffusion of both attractant and autoattractant was increased together, the amount of migration resembled that of the baseline, but the parameter space of optimal migration was shifted toward shorter autoattractant lifetimes — similar to the effect seen with broken down autoattractant. Conversely, when both were decreased the area of optimal migration shifted towards longer lifetimes.

**Figure 6:**
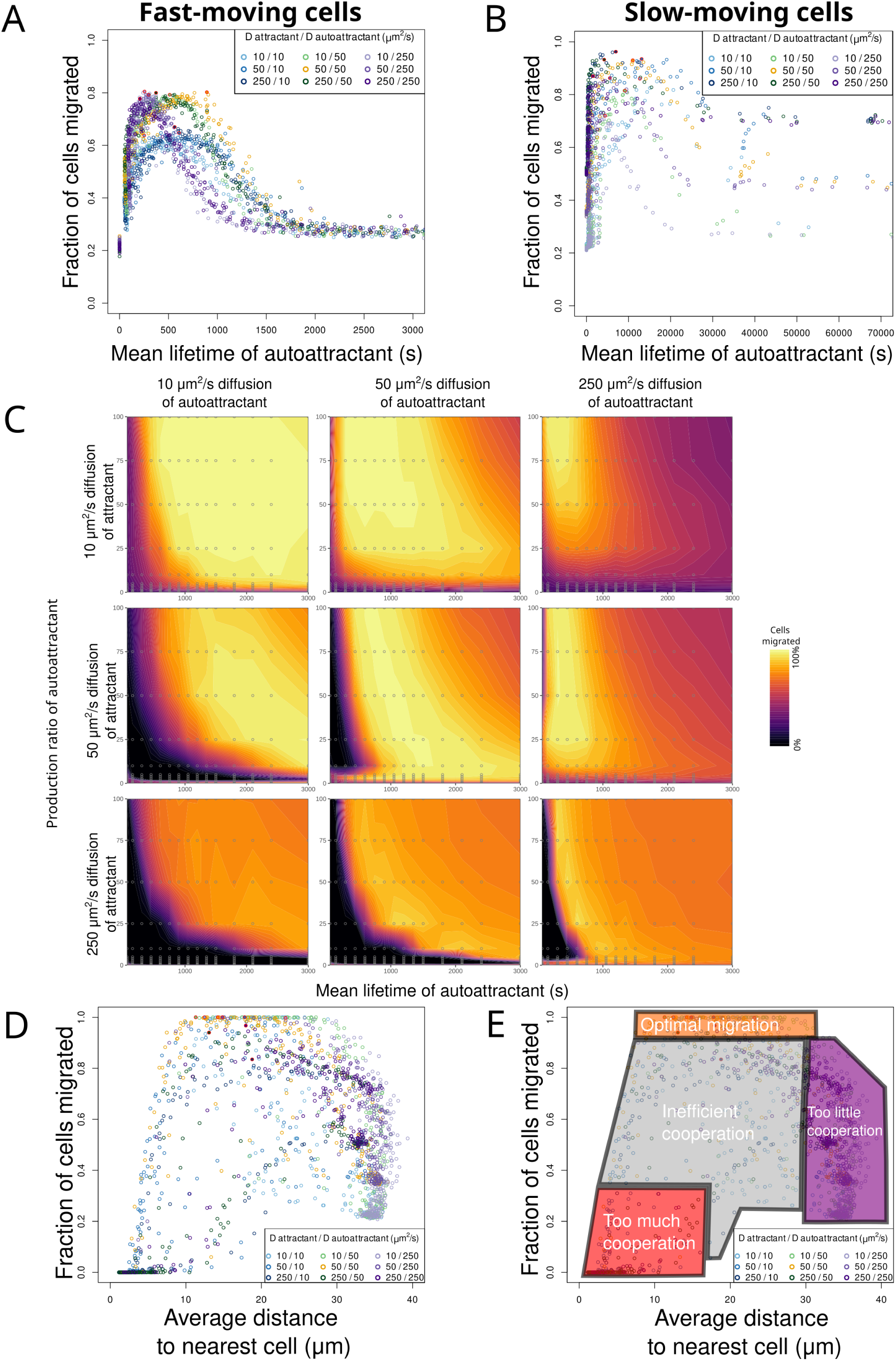
Higher diffusion promotes shorter lifetimes and efficient chemotaxis, but very short lifetimes can promote clumping. A,B. Lifetime for fast-moving (A) and slow-moving (B) cells with different diffusion speeds. Filled circles represent the best migration for that condition. *n* = 10 per point. **C.** The relationship between production and mean lifetime of a decaying autoattractant, for different diffusion speeds of attractant and autoattractant. **D.** The relation between how many cells have migrated by the end of the simulation, and their average distance to the nearest cell in the first half of the simulation, for all simulations of C. **E.** Schematic of the relation between cell migration and cooperation, based on D.

Under the conditions with high diffusion, we noted that the parameters that led to optimal migration were located very close to those that caused slow-moving clumps. To further examine this relation, we calculated the mean distance to the nearest cell for each cell in our simulations, and its relation to migration (Fig. 6D). We found that very close cells were associated with inefficient migration, but also that slightly less close cells were associated with optimal migration. Distant cells were associated with suboptimal migration. We also find that the cell-cell distance is determined by the strengths of cell- perceivable gradients across the narrow axis of the model area (Fig. S5A, calculation details in Methods section 4.5). Interestingly, the association between cell-cell distance and migration efficiency does not hold for faster-moving cells, or for slow-moving cells that break down autoattractant (Fig. S5B), but there is a clear association between the strengths of perceived gradients and migration efficiency that does hold for fast-moving cells with decaying autoattractants (Fig. S5C). Here perceptible gradients of intermediate strength are optimal, with gradients that are too weak or too strong leading to less migration. In contrast, in cells that do break down autoattractant directly there is a simple positive linear relationship between cell-cell gradient strength and migration efficiency (Fig. S5D). Taken together, this suggests that there is an optimal amount of cooperation for cells that do not break down autoattractant directly, schematically shown in Fig. 6E. An intermediate level of cell-cell attraction through autoattractants leads to optimal migration, with lower levels causing insufficient coordination between the cells near to the attractant gradient and those further away, and so leading to cells being left behind. Higher cooperation levels caused slow-moving clumps as cells coordinated too strongly with each other and too little to the primary attractant. In contrast, cells that break down autoattractant around them migrate best when they generate a very strong gradient.

## 3 Discussion

Our model makes several key predictions about the role of autoattractants in collective cell migration. Firstly, autoattractants enhance collective migration by providing a secondary signalling layer that allows trailing cells to follow the path of frontrunner cells, even when those trailing cells cannot directly sense the primary attractant gradient. This is consistent with the signalling observed in neutrophil swarming [21, 30] and *Dictyostelium* aggregation [27].

Secondly, autoattractants must be short-lived to be functional, as a long-lived autoattractant cannot give useful information on the location of the front of a moving wave. Optimal autoattractant lifetimes depend on cell and attractant characteristics more than on the environment. While the specific production and breakdown rates that yield optimal migration vary with cell density and primary attractant concentration, the optimal lifetime for autoattractants is remarkably consistent. This suggests that many autoattractants may have tightly controlled short lifetimes.

Thirdly, decaying autoattractants those removed by spontaneous degradation or ubiquitous enzymes rather than by cellular breakdown are more robust across environments, although they achieve a somewhat lower migration optimum. This robustness arises because the autoattractant lifetime is determined by a fixed decay rate rather than by the variable number of consuming cells, so that the lifetime varies much less across conditions.

Fourthly, slow cells are generally more efficient at migration relative to their speed, but can also clump more easily. This clumping occurs when the front cells are attracted more towards each other than towards the primary attractant, and so stop moving directionally. We find that efficient migration relies on close cell-cell interactions that are not far removed from conditions that lead to clumping instead of migration.

### 3.1 Implications for *in vivo* systems

The dynamics we describe are likely prevalent in many more systems. Autoattractant- like behaviours have been most extensively studied in *Dictyostelium*, where cAMP relay drives collective aggregation [27], and in neutrophils [1, 30], but are also key in microglia [10, 8]. We have shown that close associations of cells that communicate through short-lived autoattractants are the most efficient at collective migration. Because they would be short-lived, it is inherently difficult to find additional autoattractants *in vitro* or *in vivo*: samples collected would be depleted of autoattractant before measurements could be made. New techniques may find autoattractants in many more cell types.

In this text we have assumed that the attractant and autoattractant receptors have the same Kd, but this is not a requirement of the model. For example, an autoattractant that was produced with a tenfold higher production ratio and a tenfold higher Kd, Km and Vmax would show identical behaviour (eqs. 5,10,11).

We have described good migration here as "optimal", but it is important to keep in mind that recruiting of a large number of cells is one of several outcomes that may be selected for *in vivo*. In many biological systems, evolution will have shaped cells to exhibit a particular level of migration, which will often be only limited migration rather than maximal migration. This is reflected in the diversity of existing systems that may use autoattractants - each likely tuned to its specific functional requirements. Similarly, the clumping behaviour we observe could be an advantageous effect of autoattractant signalling in contexts where it is desirable for cells to find each other while migrating. For example, in lymph nodes, the aggregation of dendritic cells and T cells is functionally important for initiating adaptive immune responses, and autoattractant-driven clustering could facilitate this process. [12, 5]

Autoattractants released in response to primary attractants, as modelled here, should be distinguished from migration with co-attraction, where cells constitutively attract each other regardless of external gradients [23, 9]. In our model, autoattractant production depends directly on sensing the primary attractant, creating a signal that transmits information about the direction of migration, rather than a simple aggregation signal. However, as we have shown, the signal can also become primarily an aggregation signal when it is so short-lived that it forms very strong local gradients.

### 3.2 Future extensions

Several extensions of this model merit future investigation. First, further enhancement of migration may exist through the production of an inhibitor derived from the autoattractant. Such an inhibitor could bind the same receptors as another attractant, but not provide as strong of a chemotactic signal, and so could create a smaller but more directional secondary wave [11]. Variation in chemotactic response between different receptors in general would also allow the model to potentially cover more situations. Second, pulsatile release of autoattractant, as observed in neutrophil swarming [30], could modulate the temporal dynamics of the signal. Finally, relay of the autoattractant signal, where cells amplify the signal by producing additional autoattractant in response to sensing it, could extend the effective communication range, as observed in microglial ATP signalling [10].

## 4 Methods

### 4.1 Model overview

We developed a multi-scale agent-based model of immune cell chemotaxis in which individual cells are represented as off-lattice circular agents operating on a two-dimensional regular lattice of chemical concentrations. The simulation represents a chemotaxis chamber with cells initially placed at one end and a primary attractant uniformly distributed throughout the domain. The model integrates chemical diffusion dynamics with discrete cell decision-making at each timestep.

### 4.2 Environment and chemical dynamics

#### 4.2.1 Lattice and geometry

The simulation domain is a two-dimensional grid of size *N_x_ × N_y_* lattice sites, where each site represents a Δ*x* = 10 *µ*m square. The domain is bounded by no-flux boundary conditions. Two chemical species are tracked on the lattice: a primary attractant and an autoattractant, each stored as a concentration value at every lattice site. In what follows, *C_i,j_* denotes the concentration at site (*i, j*) of a generic species (each species obeys the dynamics below independently); where species must be distinguished (Eq. 5) we write *C_ℓ,i,j_* for ligand *ℓ*.

#### 4.2.2 Diffusion

Chemical diffusion is computed each timestep using the DuFort–Frankel finite difference scheme. For an interior lattice site (*i, j*) that is not a wall, the concentration *C_i,j_*is updated as:

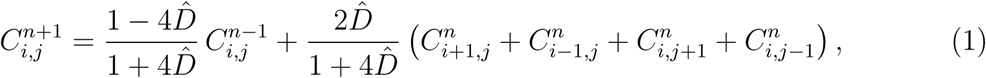

where *D*^^^ = *D* Δ*t/*Δ*x*^2^ is the dimensionless diffusion number, *D* is the diffusion coefficient (*µ*m^2^/s), Δ*t* = 6 s is the timestep duration, and Δ*x* = 10 *µ*m is the lattice spacing. If a neighbouring site is a wall, the concentration at the current site from the previous timestep is used in place of that neighbour. The first diffusion step of each timestep uses a forward Euler step to provide the two-level initial condition required by the DuFort–Frankel scheme:

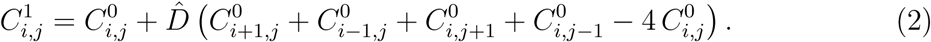

To reduce errors, multiple sub-steps of diffusion are performed per timestep, with proportionally reduced *D*^^^, so that *D*^^^ is never larger than 0.1. The lattice sites at the bottom three rows are set to a concentration of attractant equal to that of the initial condition (Table 1) at the start of every timestep to represent inflow from the environment.

#### 4.2.3 Decay of autoattractant

For the simulations with a decaying autoattractant, concentration decays at every lattice site each timestep:

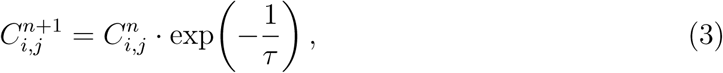

where *τ* is the decay time constant parameter (in timesteps).

### 4.3 Cell behaviour

#### 4.3.1 Cell geometry

Each cell is represented as a circle of radius *r* defining the set of lattice sites it occupies. The occupied sites *S*_cell_ are all lattice sites (defined by their midpoint (*i, j*)) satisfying:

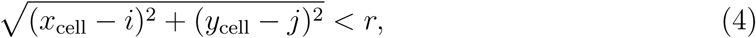

where (*x*_cell_*, y*_cell_) is the continuous-valued cell centre position.

#### 4.3.2 Gradient sensing and movement

Every timestep each cell senses the local chemical gradient across its occupied sites using a receptor-binding model with saturable kinetics. For each occupied lattice site (*i, j*) and each ligand *ℓ*, the perceived signal is computed as:

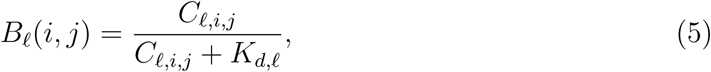

where *C_ℓ,i,j_* is the concentration of ligand *ℓ* at site (*i, j*) and *K_d,ℓ_* is the corresponding receptor dissociation constant. The net attraction *A*(*i, j*) at each site is the sum over all ligand contributions:

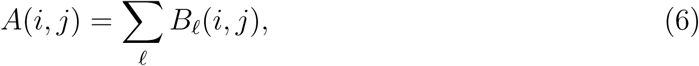

Additionally, environmental noise is added to the attractant grids before sensing by adding a uniform random value *U* (0*, σ_e_*) to every lattice site, where *σ_e_* is the environmental noise amplitude. This random value is determined separately for each lattice site and each timestep, and is not carried forward.

The target direction is then computed as the angle from the unweighted centroid of its occupied sites to the attraction-weighted centroid of its occupied sites:

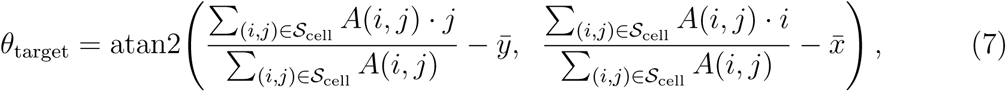

where *x̄* and *ȳ* are the arithmetic mean coordinates of the occupied sites.

The cell’s actual heading for movement is a weighted combination of its previous heading *θ*_prev_ and the newly sensed target *θ*_target_, governed by a persistence parameter *p ∈* [0, 1] which imposes an implicit limit on how sharply a cell can turn in a single timestep:

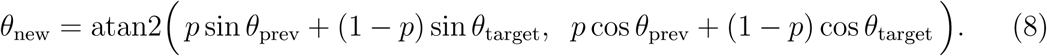

We used a persistence of p = 0.9 throughout, corresponding to the minimum time to reverse direction (i.e. turn 180 degrees) of 2.8 minutes listed in Table 2.

The cell then attempts to move a fixed distance *d* in the direction *θ*_new_:

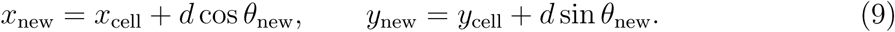

Movement is rejected if it would place any part of the cell outside the domain boundaries, cause overlap with a wall, or cause overlap with another cell. If the move is rejected the cell selects a new random heading, but will not try to move until the next timestep, which effectively slows it down. If movement is successful the cell’s lattice sites (Eq. 4) are recalculated.

#### 4.3.3 Attractant breakdown

Each cell breaks down the primary attractant, and where relevant also the autoattractant, from the lattice sites it covers using Michaelis–Menten kinetics. The amount broken down at each occupied site (*i, j*) *∈ S*_cell_ is

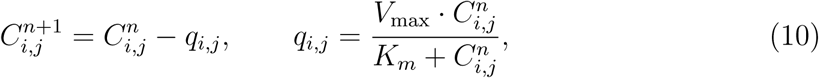

where *V*_max_ is the maximum breakdown rate and *K_m_* is the Michaelis constant. The concentration is set to zero if it would become negative.

#### 4.3.4 Autoattractant production

Each cell produces autoattractant in proportion to the total primary attractant it broke down this timestep (Eq. 10), distributed uniformly across its occupied sites. The autoattractant concentration at each occupied site (*i, j*) *∈ S*_cell_ is incremented by

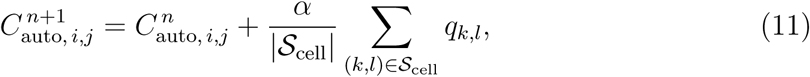

where *α* is the production ratio, *q_k,l_* is the primary attractant broken down at site (*k, l*) (Eq. 10), and *|S*_cell_*|* is the number of occupied sites.

### 4.4 Simulation parameters

Simulations were parameterised to represent two broad classes of immune cells: fast-moving cells (e.g. neutrophils) and slow-moving cells (e.g. macrophages). Environmental and cell-specific parameters are summarised in Tables 1 and 2, respectively.

For each parameter combination, *n* = 10 replicate simulations were performed with different random seeds.

### 4.5 Analysis

#### 4.5.1 Migration outcome

Migration was quantified as the fraction of cells that had crossed into the lower quarter of the simulation domain (i.e. *y < N_y_/*4) by the final timestep. For each parameter combination, *n* = 10 replicate simulations were performed with different random seeds, and the mean fraction of migrated cells was computed across replicates.

#### 4.5.2 Autoattractant lifetime

In some figures we calculate the mean effective lifetimes of autoattractant. At each recorded timestep, the total autoattractant concentration across all lattice sites was divided by the total amount of autoattractant removed by breakdown in that interval. This ratio gives the average time that a unit of autoattractant persists in the system at each point in the simulation. The mean over all recorded timesteps was then taken as the effective lifetime for that simulation.

#### 4.5.3 Gradient strength

To quantify the perceived cell-cell gradient strength, the concentration *C_i,j_* at each lattice site is first transformed using the same receptor-binding equation used for gradient sensing (Eq. 5), giving a receptor-occupancy field *B*(*i, j, t*). The gradient strength is then the summed absolute difference in receptor occupancy between neighbouring lattice sites along the short axis of the domain, normalised by the lattice size:

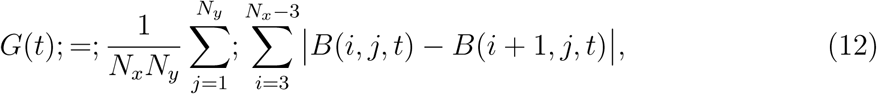

where lattice columns with *i <* 3 or *i > Nx −* 3 are excluded to avoid boundary artefacts. This yields *G*(*t*) at each recorded timestep.

### 4.6 Implementation and Data availability

The model was implemented in Python 3, and the analysis was performed in R. Simulation code is available at https://github.com/DMvers/AutoTaxis

## Acknowledgements

This work was supported by the Wellcome Trust (https://wellcome.org; grant 221786/Z/20/Z to RHI). The authors acknowledge the use of the UCL Myriad High Performance Computing Facility (Myriad@UCL), and associated support services, in the completion of this work.

## A Supplemental Figures

**Supplemental figure S1.**
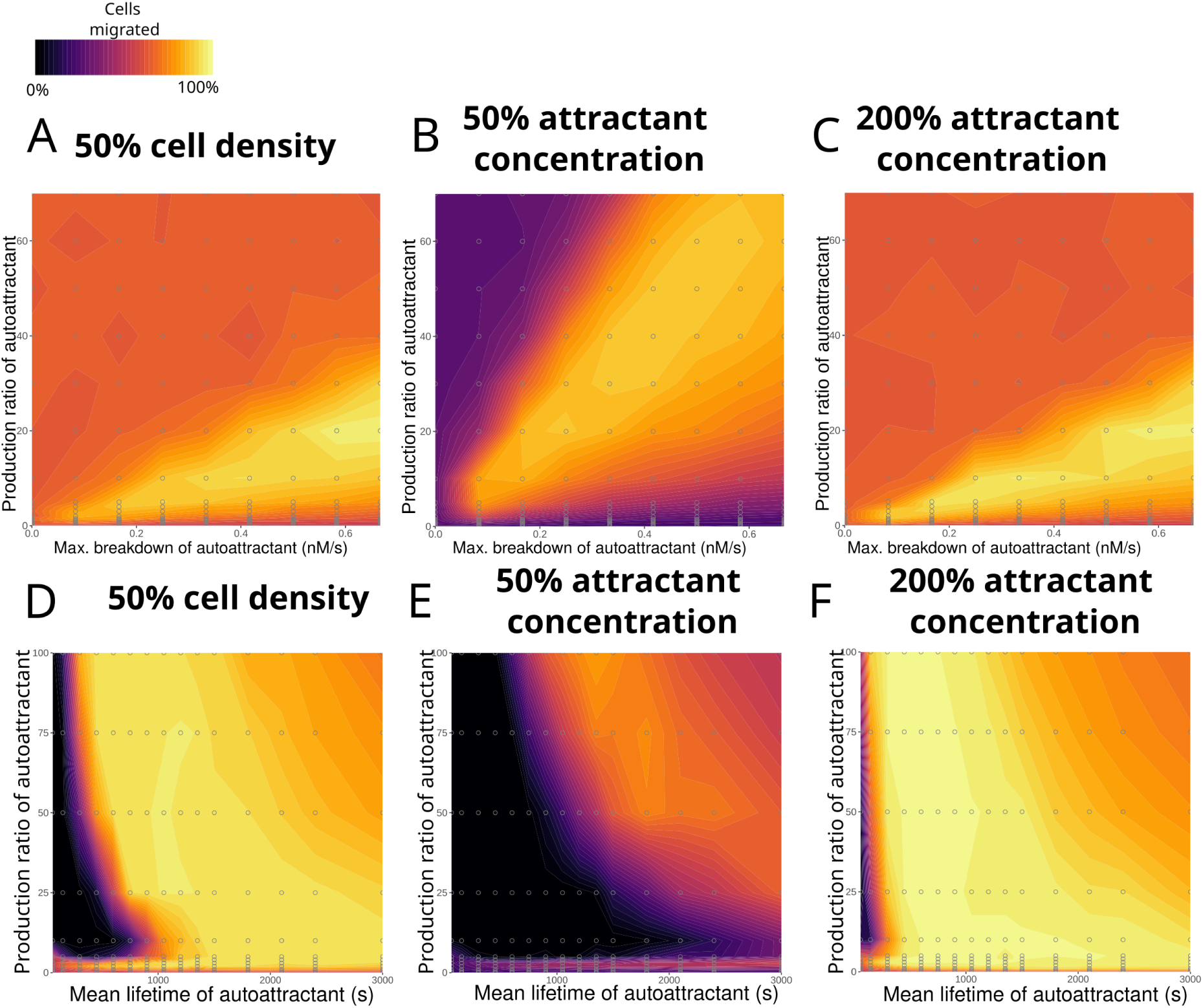
The relationship, for slow-moving cells, between autoattractant decay and production and migration outcomes for (A–C) a broken down autoattractant at low cell density, low initial primary attractant concentration, and high initial primary attractant concentration, and (D–F) the same for a decaying autoattractant.

**Supplemental figure S2.**
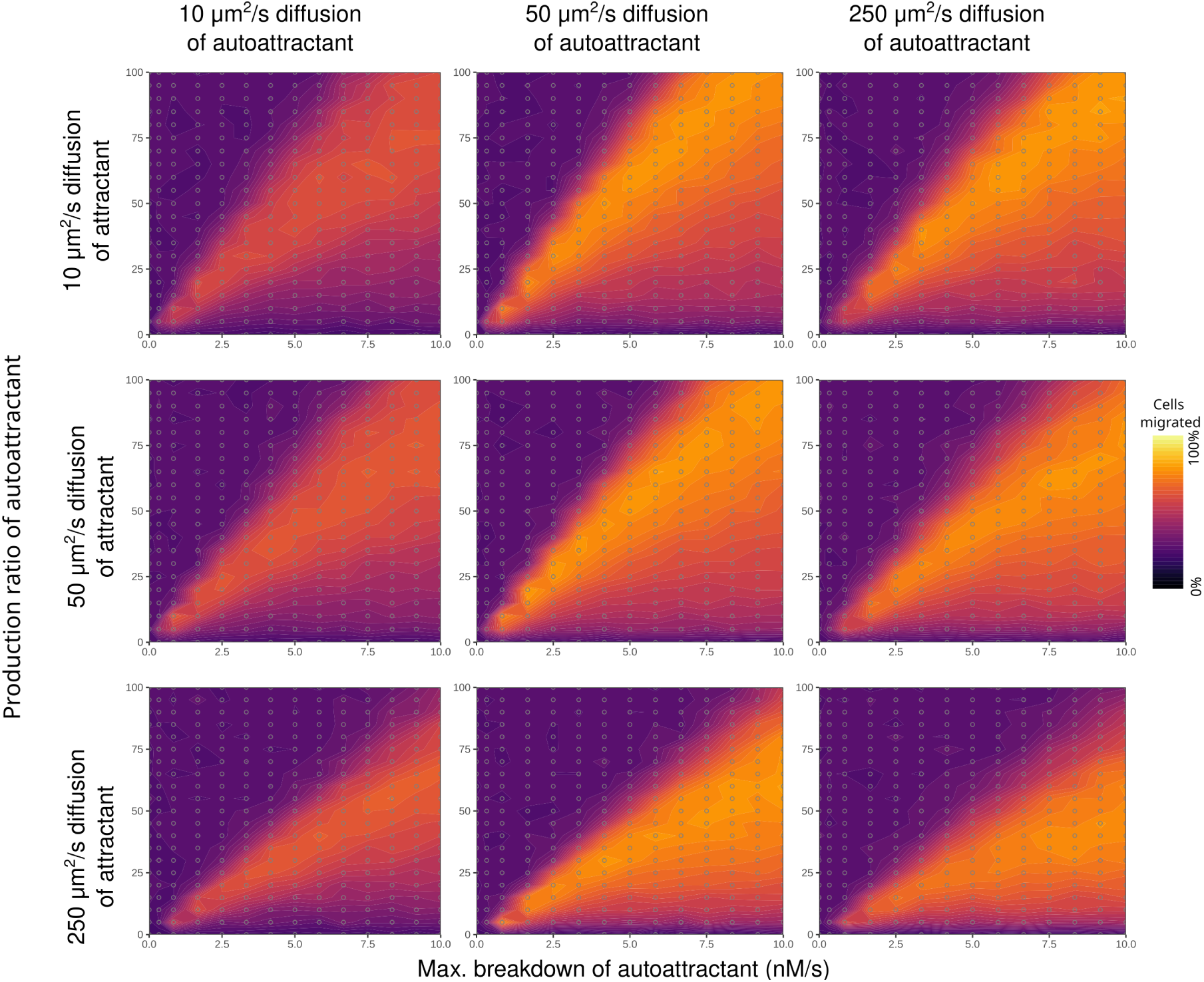
The relationship between production and breakdown for fast-moving cells and a broken down autoattractant, for different diffusion speeds of attractant and autoattractant.

**Supplemental figure S3.**
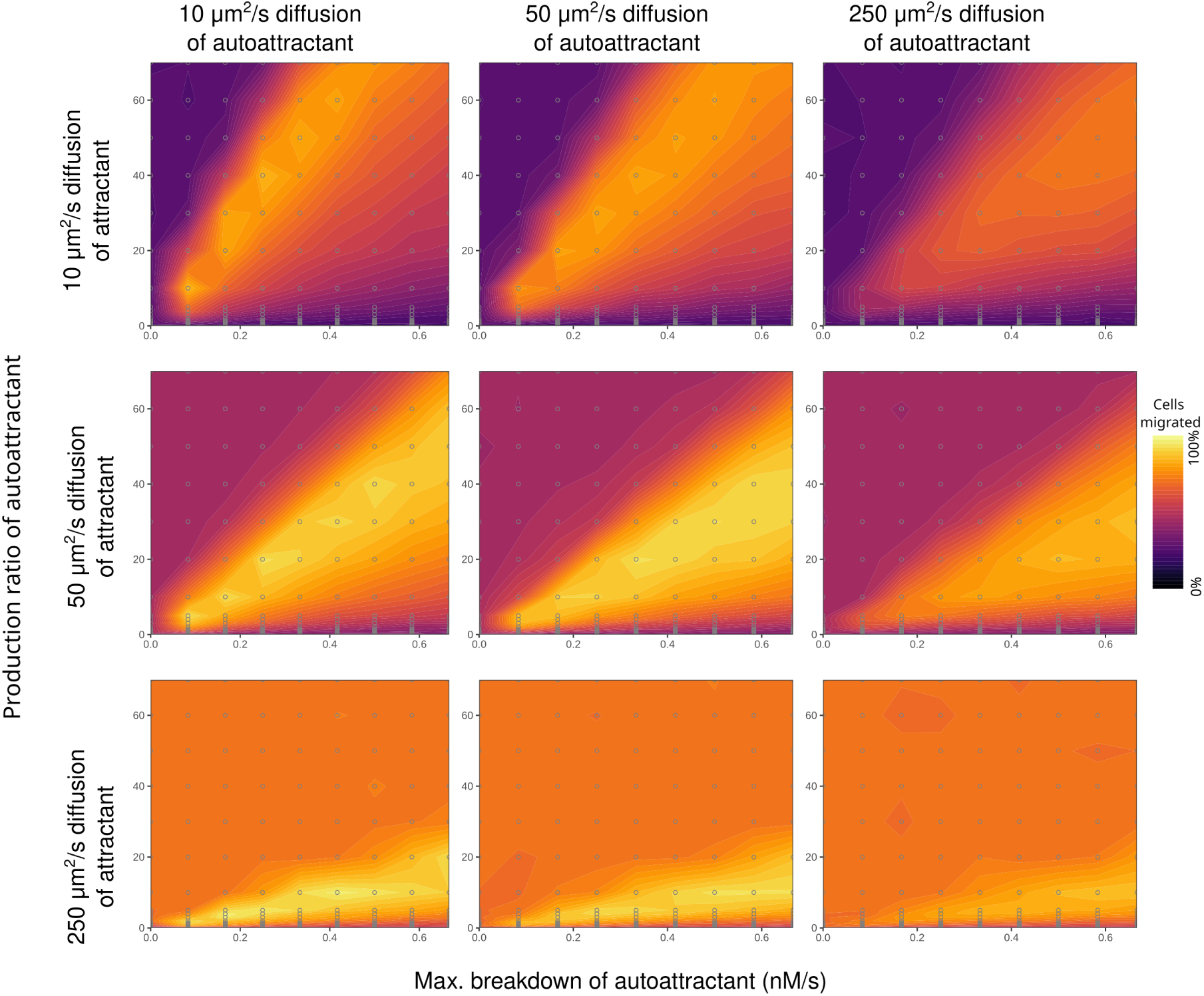
The relationship between production and breakdown for slow-moving cells and a broken down autoattractant, for different diffusion speeds of attractant and autoattractant.

**Supplemental figure S4.**
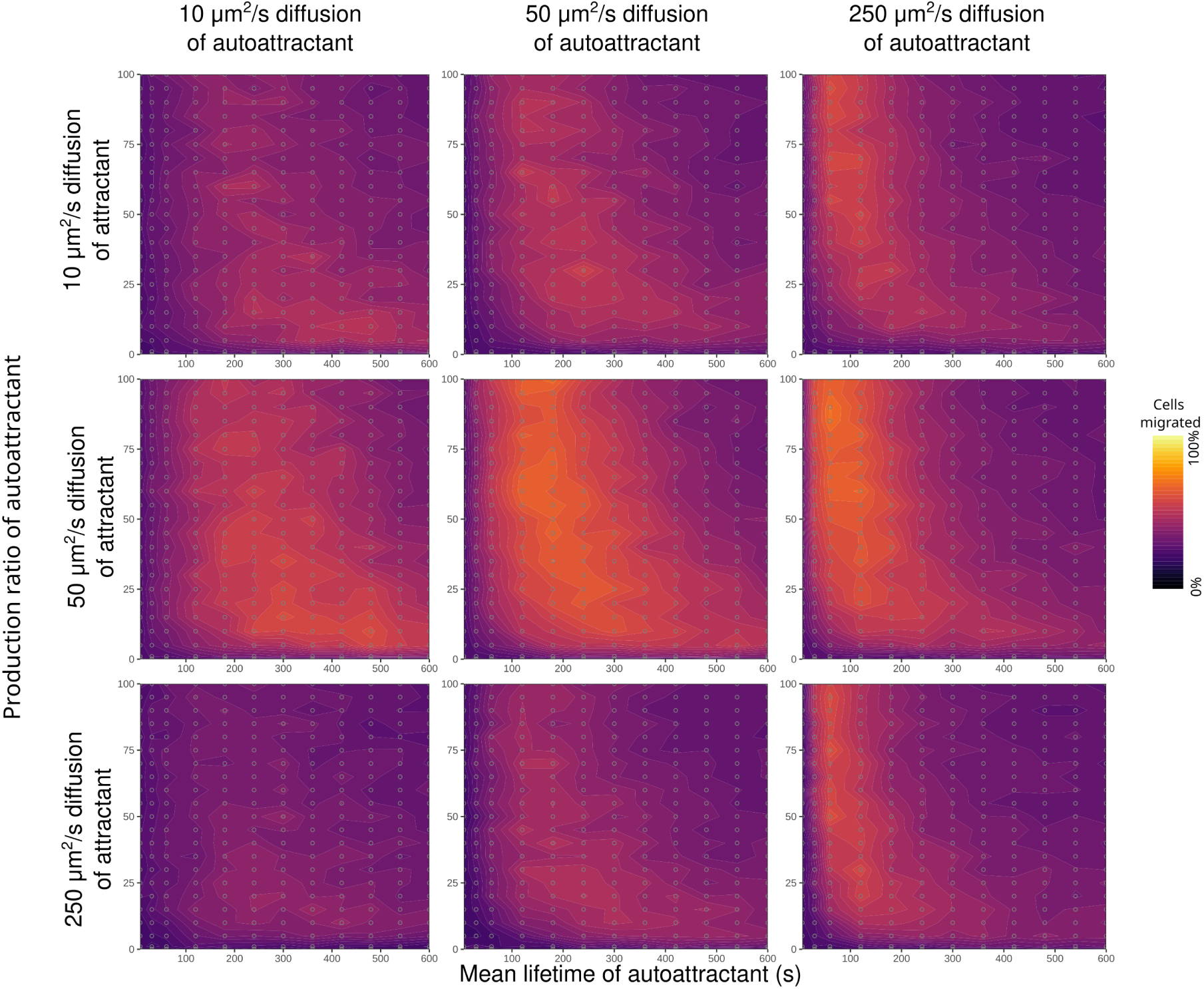
The relationship between production and mean lifetime for fast-moving cells and a decaying autoattractant, for different diffusion speeds of attractant and autoattractant.

**Supplemental figure S5.**
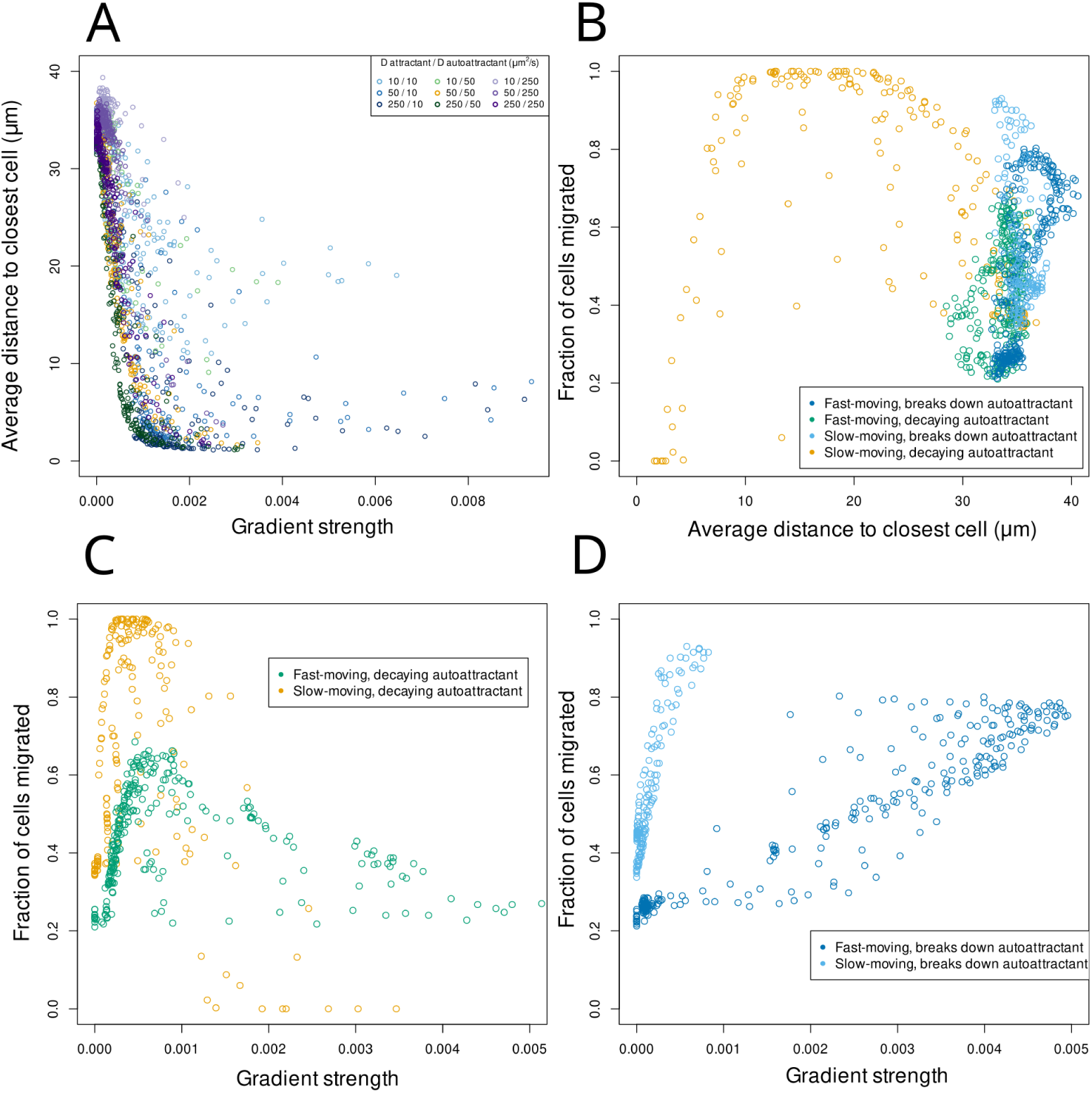
For A,C,D: Gradient strength is represented by the average difference in autoattractant receptor binding over every 10 *µ*m (one grid site) distance across the narrow side of the environment, so that . Higher gradient strength indicates stronger cell-cell attraction. Details of calculation in methods section 4.5. **A.** The relation between gradient strength and the average distance between cells in the first half of the simulation as in Fig. 6 D. **B.** The relation between the fraction of cells migrated by the end of the simulation, and the average distance between cells in the first half of the simulation, for the simulations of Fig. 3A, Fig. 4B, and Fig. 5C&E **C.** The relation between gradient strength and the fraction of cells migrated, for cells with decaying autoattractant, as in Fig. 4B and Fig. 5E. **D.** The relation between gradient strength and the fraction of cells migrated, for fast and slow-moving cells with broken down autoattractant, as in Fig. 3A and Fig. 5C.

## B Supplemental Videos

**Video S1**

Visualisation of the simulations corresponding to Fig. 2 A,B,D&E, showing the effect of different production ratios on the migration of fast-moving cells that break down autoattractant.

**Video S2**

Visualisation of the simulation corresponding to Fig. 4 A and two additional conditions from Fig. 4B, showing the effect of a decaying autoattractant on the migration of fast-moving cells that do not break down autoattractant.

**Video S3**

Visualisation of the simulations corresponding to Fig. 5 A&B, showing the effect of an autoattractant on the migration of slow-moving cells that break down autoattractant.

**Video S4**

Visualisation of the simulations corresponding to the four insets of Fig. 5 E, showing the effect of different production ratios and autoattractant lifetimes on slow-moving cells that do not break down autoattractant.

